# More is less: increased processing of unwanted memories facilitates forgetting

**DOI:** 10.1101/286534

**Authors:** Tracy H. Wang, Katerina Placek, Jarrod A. Lewis-Peacock

**Author notes:** Corresponding Author: Tracy H. Wang, Department of Psychology, University of Texas at Austin, Austin, TX 78712.

## Abstract

The intention to forget can produce long-lasting effects. This ability has been linked to suppression of both rehearsal and retrieval of unwanted memories – processes that are mediated by prefrontal cortex and hippocampus. Here, we describe an alternative account of deliberate forgetting in which the intention to forget is associated with increased engagement with the unwanted information. We used pattern classifiers to decode functional magnetic resonance imaging (fMRI) data from a task in which participants viewed a series of pictures and were instructed to remember or forget each one. Pictures followed by a forget instruction elicited higher levels of processing in ventral temporal cortex compared to those followed by a remember instruction. This boost in processing led to more forgetting, particularly for items that showed moderate (vs. weak or strong) activation. This result is consistent with the non-monotonic plasticity hypothesis, which predicts weakening and forgetting of memories that are moderately activated.

---

The human brain is not capable of remembering everything – in our lifetime we will forget the majority of our experiences. While this may seem a bleak consequence, memory loss is essential to the human experience; we are bombarded with too much information each moment to possibly record and preserve every experience. Which information should be saved and which should be discarded? This challenge is often solved automatically by the brain (e.g., through automatic learning processes such as statistical learning^1^). However, there are instances in which people have volitional control over what they will expunge from their memory^2,3,4^. In these cases, forgetting can be considered an adaptive feature of memory in which unwanted or irrelevant information is actively discarded to improve access to other memories.

This ability to intentionally forget an unwanted experience has been shown to involve inhibitory processes in frontal control regions that act to suppress the undesired information. Attempts to forget new memories have been linked to increased activity in right dorsolateral prefrontal cortex and decreased activity in hippocampus^5.6^. Furthermore, these studies also show that an increase in the functional coupling between these two regions leads to successful intentional forgetting. It is unclear, however, how sensory representations of the ‘to-be-forgotten’ memories in posterior cortical areas^7,8,9^ are related to the success of deliberate forgetting.

In the present experiment, we hypothesized that deliberate forgetting is facilitated by the weakening of moderately active memories in sensory cortex. This idea is motivated by the non-monotonic plasticity hypothesis^10,11^ which states that moderately active memory representations are weakened by local inhibitory mechanisms in brain regions supporting their representation^12^. Our prior work demonstrated that non-monotonic plasticity contributes to incidental forgetting of items in working memory^13^. When participants did not decisively switch their attentional focus between two items in working memory, the previous item would linger in a state of moderate activation (as measured by pattern classifiers applied to fMRI data). According to the non-monotonic plasticity hypothesis, these items were susceptible to weakening, and indeed we found that they were associated with worse subsequent memory, compared to trials with less lingering activation. Here, we sought to test whether the weakening of moderate memory activations might also play a role in deliberate forgetting. The non-monotonic plasticity hypothesis predicts that ‘to-be-forgotten’ items that have a moderate degree of activation during encoding should be associated with worse subsequent memory relative to items for which there is greater or lesser activation.

To test this prediction, we used an item-method directed forgetting paradigm^3,4^ in which participants were presented with pictures of male/female faces and indoor/outdoor scenes, one at a time, with each picture followed by an instruction either to remember that picture for later, or to try and forget having ever seen it (Fig. 1c). All pictures, regardless of memory instruction, were later presented along with novel pictures of faces and scenes in a recognition memory test at the end of experiment. No specific instructions were given to participants regarding *how* to remember or forget any of the pictures. We expected that participants would respond to a remember instruction by maintaining their focus on that item to strengthen its encoding via rehearsal or elaboration. We hypothesized that participants would respond to a forget instruction by changing the amount of attention directed towards that item, and hence to alter its state of memory activation^14,15,16^. If participants withdrew attention from a ‘to-be-forgotten’ item, this would decrease that item’s activation relative to ‘to-be-remembered’ items. Alternatively, if participants increased their attentional focus on a ‘to-be-forgotten’ item, this would increase memory activation of that item. To quantify and track memory activation, we applied pattern classifiers to human fMRI data to measure processing of face and scene items throughout each trial. We focused our classifiers on activity in ventral temporal cortex, a region that serves as input to convergence zones (for example, in the medial temporal lobes) that are responsible for storing long-term memories^17^; this allowed us to treat our scene and face classifier evidence scores as reflecting the strength of the excitatory inputs into memory regions. We then related these neural measurements of memory activation (classifier evidence) to subsequent memory performance on the recognition test for these items at the end of the experiment. We found that ‘to-be-forgotten’ items were associated with worse subsequent memory overall and *higher* post-instruction classifier evidence, on average, compared with ‘to-be-remembered’ items. Moreover, the degree of activation for ‘to-be-forgotten’ items was predictive of their subsequent memory strength. This result suggests a new perspective on intentional forgetting in which, when trying to forget memories, people raise the amount of their lingering activity in order to weaken them.

**Figure 1.**
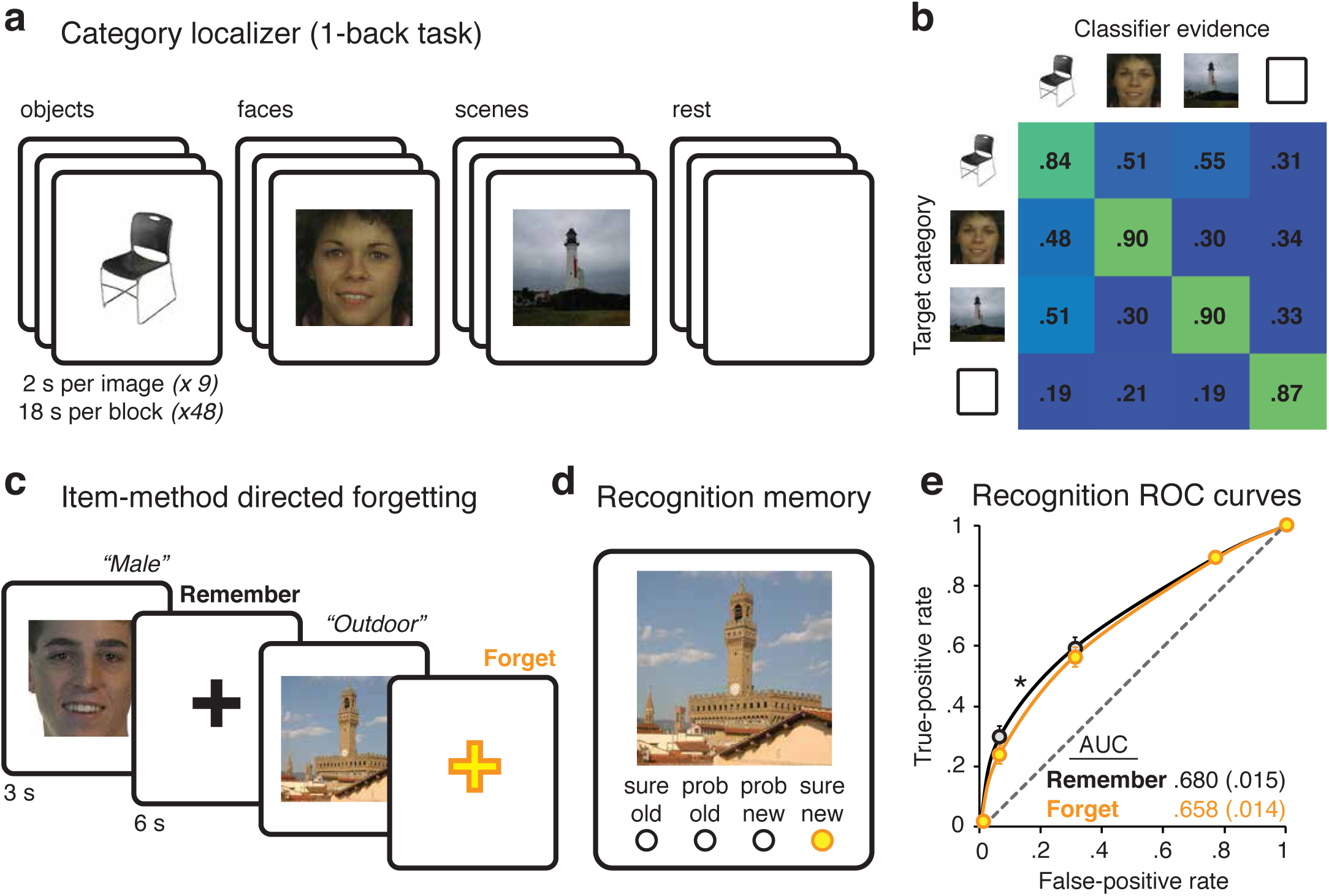
Task procedures, classifier sensitivity, and subsequent memory performance. **(a)** Participants performed a category localizer (1-back task) in the scanner with objects, faces, scenes, and rest. **(b)** Classifier evidence scores (between 0 and 1) for each target category were obtained from cross-validation analysis of fMRI data from the localizer. **(c)** Next participants performed an item-method directed-forgetting task on faces and scenes in the scanner. They made a subcategory judgment on each picture, and then a cue appeared telling them either to remember (black cross) or to forget (yellow cross) that picture. **(d)** At the end of the experiment, participants were given a recognition memory test for faces and scenes that were studied, and for new faces and scenes. **(e)** Recognition memory sensitivity was assessed by receiver operating characteristic (ROC) analysis separately for Remember and Forget items. AUC, area under the ROC curve. * *P* = 0.021. Error bars indicate the s.e.m, *n*=20.

## RESULTS

### Behavioral results

Figure 1a and 1c shows the design of the experiment. Participants performed a category localizer task in the scanner. fMRI data from this task were used to train a category-level classifier to distinguish processing of objects, faces, scenes, and rest in ventral temporal cortex. Participants then performed an item-method directed forgetting task on pictures of faces and scenes. On each trial, a stimulus appeared and participants made a subcategory judgment (male/female; indoor/outdoor), and then they received an instruction to remember or to forget that picture. No specific strategies were provided to participants. At the end of the experiment, participants were given a (behavioral) recognition test for the pictures that had been studied in the directed forgetting task.

The perceptual localizer consisted of a 1-back task on miniblocks of same-category items (i.e. face, scene, object and rest) for category-level decoding. We obtained accuracy (successfully identifying a repeated image) and response latency behavioral performance measures. Outlier trials for which response latencies were greater than 3 standard deviations from each subject’s mean were removed from the analysis (2.2% of all trials). Accuracy on the 1-back task was at ceiling for faces (97.7%, 0.5%s.e.m.), scenes (98.2%, s.e.m. 0.4%), and objects (98.5%, s.e.m. 0.4%). Rest trials did not require a response. One-way analysis of variance (ANOVA) revealed no accuracy differences (*P* =.530) between these three categories. Further, one-way ANOVA test of reaction times across faces (593ms, s.e.m. 25ms), scenes (614ms, s.e.m. 33ms) and objects (600ms, s.e.m 28ms) revealed no differences between these categories (*P* =.871). Subcategory identification accuracy in the directed-forgetting task was high for both faces (97.7%, s.e.m. 0.4%) and scenes (98.0%, s.e.m. 0.4%), with no significant differences between them (*P* =.570 two-tailed paired t-test). Participants responded well within the 3-sec response deadline, and were faster to identify faces (1061 ms, s.e.m. 42 ms) than scenes (1236 ms, s.e.m. 45 ms; *P* <.001, two-tailed paired t-test). Performance on the subsequent memory test of these items is shown in Fig. 1d. For all subsequent memory analyses of old items described below, we treated recognition responses as a graded measure of memory strength (sure old = 1, probably old = 0.667, probably new = 0.333, sure new = 0), in which the ‘old’ responses corresponded to remembered items and ‘new’ responses corresponded to forgotten items. Recognition memory sensitivity was significantly above chance for both ‘to-be-remembered’ items (two-tailed *t*-test on area under the receiver operating characteristic curve, *t*(19) = 11.89, *P* <.001), and for ‘to-be-forgotten’ items (*t*(19) = 11.33, *P* <.001). Participants showed successful intentional forgetting^7^ as memory performance was significantly worse for ‘to-be-forgotten’ items (*t*(19) = 2.52, *P* = 0.021 two-tailed paired *t*-test).

### Neural measures of directed forgetting

We conducted univariate fMRI analyses based on the general linear model to contrast brain regions engaged for ‘to-be-forgotten’ items that were forgotten and ‘to-be-remembered’ items that were remembered (see Methods). Consistent with prior work^5,18,19^, we found increased activity for successful forgetting in dorsolateral prefrontal cortex (DLPFC), left ventral medial prefrontal cortex (VMPFC), posterior cingulate, and precuneus (Supplementary Fig. 1) For successful remembering, we found increased activity in bilateral hippocampus and left ventral inferior frontal cortex. For a detailed report of the univariate results, please refer to Supplemental Table 1. Together with the behavioral results reported above, these data confirm that our experiment is producing directed-forgetting results that are consistent with prior findings.

### Measuring memory processing with fMRI pattern classifiers

To assess the degree of memory processing on each trial, we applied pattern classifiers to the fMRI data^7,8,20^. Group-averaged results for the classifiers, trained separately for each participant’s localizer data, are shown in Fig. 1b. The classifier confusion matrix shows the mean classifier evidence for all categories (columns) on localizer blocks featuring stimuli from one target category (rows). The cross-validation procedure used to evaluate classifier performance entailed training a classifier on two runs of data and then applying that classifier to independent data from the held out third run; the runs were then rotated and this procedure was repeated until all three runs had been tested. Decoding accuracy for all categories was well above chance (25%). Face evidence was reliably higher than scene evidence for face blocks, and vice versa (both *P*’s < 0.001), but face and scene scores were not dissociable during rest periods (*P* = 0.650). To analyze data from the memory task, we applied classifiers that were retrained on all localizer data, again separately for each participant. For each trial, we computed a “target-nontarget” classifier evidence score which reflects the relative balance between trial-relevant processing and trial-irrelevant processing (e.g., “face evidence minus scene evidence” on a face trial). These neural measures served as a proxy for item-specific processing on each trial. Averaged across trials, the classification results show that evidence for the memory item was higher after a Forget instruction compared to after a Remember instruction (between 6 and 8 s after stimulus onset, two-tailed t-test, *P* = p<.001; Fig. 2a). This indicates stronger processing of ‘to-be-forgotten’ items. Importantly, this is inconsistent with a prominent view that item-method directed forgetting results from stronger encoding (e.g., selective rehearsal) of ‘to-be-remembered’ items^21,22.^ Instead, it agrees with more recent reports that deliberate forgetting is associated with effortful processing following an instruction to forget^23-27^. Forgetting effects have been linked to a decrease in memory sensitivity for ‘to-be-forgotten’ items, as we found here, rather than outright forgetting of those items. For example, Zwissler and colleagues^28^ found that forget instructions result in active processing that reduces the false-alarm rate, but does not impair memory beyond an uncued baseline where only incidental encoding occurs. Here, our central hypothesis is that the degree of memory processing after a forget instruction will predict the degree of forgetting success for that item^10-13^. We now address this hypothesis by linking the neural measures of memory processing during directed forgetting with the behavior measures of memory sensitivity from the recognition test at the end of the experiment.

**Figure 2.**
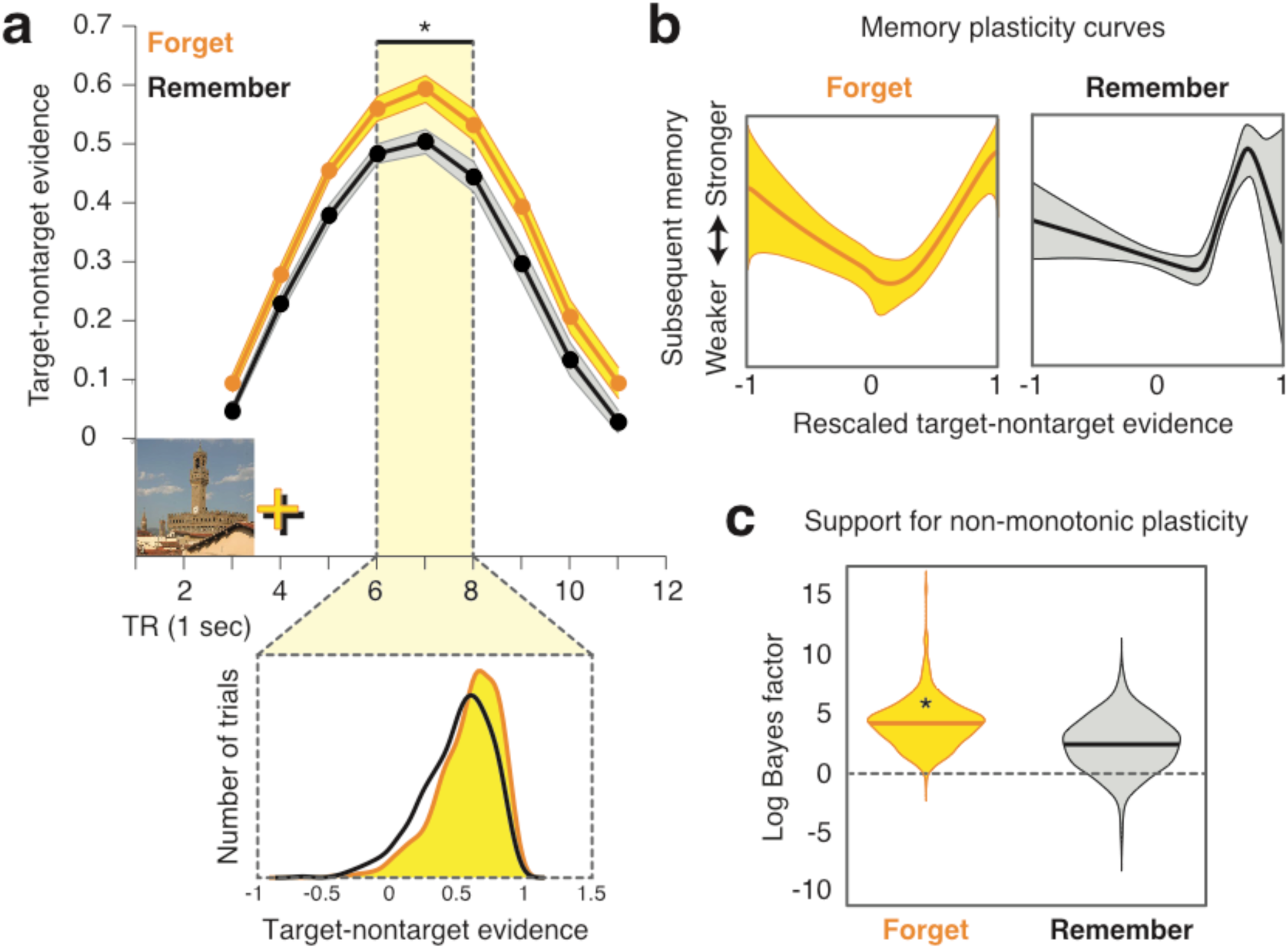
Pattern classification of fMRI data from directed-forgetting task. **(a)** Target-nontarget classifier evidence for Forget (yellow) and Remember (black) trials. Classifier evidence scores were not shifted to account for hemodynamic lag. (Ribbon thickness indicates s.e.m. across participants, *n* = 20; \**P* <.001. (bottom): Probability density distribution of the number of Forget (yellow) and Remember (black) trials based on classifier evidence scores during the peak response to each stimulus (6-8 sec post onset). **(b)** Empirically derived estimates (generated using the Bayesian P-CIT algorithm^4^) of the ‘plasticity curve’ characterizing the relationship between target-nontarget classifier evidence and subsequent memory performance (recognition confidence). Behavioral outcomes are modelled as depending on the summed effects of pre-instruction (1-3 s) and post-instruction (4-9 s) classifier evidence. Within each box, the line shows the mean of the posterior distribution over curves and the ribbon shows the 90% credible interval (such that 90% of the curve probability mass lies within the ribbon). The horizontal axis shows target-nontarget classifier evidence scores rescaled so that the minimum classifier evidence value = −1 and the maximum classifier evidence value = 1; the vertical axis represents the subsequent memory strength. **(c)** Violin plots describing the balance of evidence (operationalized in terms of log Bayes factor) in favor of the non-monotonic plasticity hypothesis, shown separately for the two conditions. These plots show the probability density (using kernel density estimation) of the log Bayes factor derived from 1,000 bootstrap iterations. The thick marker inside each plot indicates the mean. Positive values of the log Bayes factor correspond to evidence in favor of the non-monotonic plasticity hypothesis and negative values correspond to evidence against the hypothesis. (*\*P* = 0.019).

### Relating classifier evidence to subsequent memory

We hypothesized that across items, there would be a non-monotonic (U-shaped) relationship^10,11^ between target-nontarget classifier evidence for ‘to-be-forgotten’ items and subsequent recognition memory for those items at the end of the experiment. To formally test for the non-monotonic pattern in these data, we used the Probabilistic Curve Induction and Testing Toolbox (P-CIT^11,13^) Bayesian curve-fitting algorithm to estimate the shape of the ‘plasticity curve’ relating post-instruction memory processing in the directed-forgetting task (indexed by classifier evidence) and subsequent memory performance (indexed by recognition confidence). The P-CIT algorithm approximates the posterior distribution over plasticity curves (that is, which curves are most probable, given the neural and behavioral data). P-CIT generates this approximation by randomly sampling curves (piecewise-linear curves with three segments) and then assigning each curve an importance weight that quantifies how well the curve explains the observed relationship between the neural and behavioral data. Finally, these importance weights are used to compute the probability of each curve. To assess evidence for the non-monotonic plasticity hypothesis, P-CIT labels each sampled curve as theory consistent (if it shows a U shape, dropping below its starting point and then rising above its minimum value) or theory inconsistent, and then computes a log Bayes factor score that represents the log ratio of evidence for versus against the non-monotonic plasticity hypothesis; positive values of this score indicate a balance of evidence in support of non-monotonic plasticity. P-CIT also computes a χ^2^-test that assesses how well the curve explains the data overall, regardless of its shape; the P-value for this χ^2^-test indicates the probability of obtaining the observed level of predictive accuracy, under a null model where classifier evidence is unrelated to memory behavior.

For our P-CIT analyses, the pre-instruction interval and the post-instruction interval (for each item) were treated as separate learning events whose effects were summed to model recognition of that item. The fitted curves explained a significant amount of variance in subsequent recognition outcomes on both Forget trials (χ^2^ = 21.34, *P* < 0.001) and Remember trials (χ^2^ = 56.6, *P* < 0.001). Most importantly, the curves recovered by P-CIT on Forget trials revealed a U-shaped mapping between classifier evidence scores and subsequent memory outcomes, such that moderate levels of target-nontarget evidence were associated with worse subsequent memory than higher and lower levels of target-nontarget evidence (log Bayes factor = 3.90, Fig. 2c). That is, deliberate forgetting was most successful when the ‘to-be-forgotten’ item’s memory activation was sufficiently enhanced (but not too high) so as to produce moderate levels of activity during the forgetting attempt. This result held when using raw classifier evidence scores for the target category were used to quantify memory processing on each trial (e.g. “face” evidence instead of “face-scene” evidence on a face trial; see supplementary figure 2). This suggests that deliberate forgetting does not require competition per se between two or more items in memory, but rather depends on *moderate activity* of the targeted memory alone. This result is predicted by the non-monotonic plasticity hypothesis^10^, which links moderate activation with memory weakening. Note that competition between memories is one way to achieve moderate activation^13^, but it is not required^10,11, 29^.

To assess the population-level reliability of the U-shaped curve (that is, were the results driven by a small subset of participants), we also ran a bootstrap resampling test in which we resampled data from participants with replacement and re-computed the log Bayes factor for the resampled data^30^. For Forget trials, 98% of these bootstrap samples (out of 1,000 total) showed evidence in support of the non-monotonic plasticity hypothesis (that is, a positive log Bayes factor), thereby indicating a high degree of population-level reliability in the shape of the curve (Fig. 2c). There was less population-level reliability in the shape of the curve for Remember trials (only 81% of bootstrap samples showed evidence in support of a U-shaped curve).

## DISCUSSION

Here, we applied machine learning methods to human fMRI data to reveal a novel mechanism that is involved in intentional forgetting: *the weakening of moderately active representations of ‘to-be-forgotten’ items in ventral temporal cortex.* The intention to forget an item is associated with higher fMRI pattern classifier evidence, and worse subsequent memory, compared with the intention to remember an item. This boost in activation can render the memory vulnerable to disruption and therefore more susceptible to subsequent forgetting^13^. These findings are predicted by the non-monotonic plasticity hypothesis^10,11^, and they converge with recent work that describes intentional forgetting as an active and effortful cognitive process^18,19,24,26,31^. Prior work has linked forgetting to the suppression of sensory representations during the retrieval^32^ or simulation^33^ of episodic memories. We believe that the present findings provide a first step to understanding the role of modulating sensory representations during encoding to facilitate deliberate forgetting.

Our findings are compatible with and extend existing explanations for intentional forgetting^6^. In one prominent view on the neural mechanics that support intentional forgetting, Anderson and colleagues^5,6,34^ have described distinct neural mechanisms associated with two common behavioral strategies described as ‘direct suppression’ and ‘thought substitution’. Behaviorally, direct suppression is thought to be a termination of the rehearsal of items that are given a forget cue^35,36^. Direct suppression is thought to occur when inhibitory signals from dorsolateral prefrontal cortex down-regulate hippocampal engagement related to memory encoding. Thought substitution, on the other hand, occurs when subjects replace a ‘to-be-forgotten’ item with some alternative item, for example an item that was studied previously or any other random thought. During thought substitution, left ventral prefrontal cortex engages in cognitive control processes that result in demonstrative increases in hippocampal engagement^5^. In the current experiment, we did not constrain behavioral strategy to avoid any mitigation of directed-forgetting effects due to self-evaluation of instructed strategies^37^. Therefore participants may have attempted either or both thought-substitution and direct-suppression strategies, and perhaps other idiosyncratic strategies too. In spite of potentially varied strategy choices, our results show a consistent increase in memory processing following a forget instruction relative to a remember instruction. The degree of this boost in processing, specifically when it resulted in moderate activation of the item, was predictive of successful forgetting. This suggests a new route to successful forgetting: to forget a memory, its mental representation should be enhanced to trigger memory weakening (described by non-monotonic plasticity^10,11^) via local inhibitory processes governing homeostatic regulation of neural activity.

A possible limitation to this interpretation of our data comes from the categorical nature of fMRI pattern classifiers used to measure memory processing. An increase in category-specific memory processing observed for ‘to-be-forgotten’ items could result, not from an increase in processing of the ‘to-be-forgotten’ item, but perhaps from the selective retrieval and rehearsal of *another* item from the same category (e.g., rehearsing a previously studied ‘to-be-remembered’ face when instructed to forget a different face on the current trial). This would be an example of a thought-substitution strategy (described above). We argue, however, that if people were to engage in selective rehearsal of previous items, it is unlikely that they would be able to limit this process to only same-category items rather than rehearsing a mixture of previous ‘to-be-remembered’ items from both categories. The category status of a ‘to-be-forgotten’ item is an inefficient search query for previous items – the most recent same-category item would be most salient of those items in memory, and if this item had also been accompanied by a forget instruction, it would be counterproductive (for forgetting of that item) to direct attention towards it, even if to reject it, during a memory search on the current trial.

Turning to our data, we found that ‘to-be-forgotten’ items were associated with higher classifier evidence for the target category (e.g. “face” on a face trial), and also *lower* classifier evidence for the nontarget category (“scene”) relative to ‘to-be-remembered’ items (Supplemental Fig. 2). Following an instruction to forget, activation of the ‘to-be-forgotten’ item was selectively enhanced. On the other hand, there was greater evidence for task-irrelevant processing (associated with the nontarget category) after a remember instruction. This would be consistent with the idea that, following an instruction to remember, a “covert rehearsal” strategy was used in which a mixture of previously studied ‘to-be-remembered’ items (some faces, some scenes) were rehearsed^3^. Critically, if the increase in target-related processing of ‘to-be-forgotten’ items reflected rehearsal of same-category alternatives, this would predict a *linear* relationship with subsequent memory such that higher processing (indicating more same-category substitution) would lead to more forgetting. Our analyses, however, revealed a U-shaped relationship with memory processing and forgetting success, such that a moderate level of processing (but not high levels) was associated with more forgetting.

Another possible challenge to our interpretation is that an increase in category-specific information (as seen for ‘to-be-forgotten’ items) actually reflects a decrease in item-specific information^38^. That is, if the item-specific features of a representation are suppressed, leaving behind the category-generic features, this could produce higher category-specific classifier evidence scores. In order to eliminate this possibility entirely, future studies should incorporate item-level classification to determine the specificity of increased memory processing during intentional forgetting.

The strength of the current study is the identification of a relationship between lingering activation of ‘to-be-forgotten’ memories in ventral temporal cortex and their subsequent memory strength. We found that the intention to forget a memory is associated with increased processing (and neural activation) of that memory. In accordance with the non-monotonic plasticity hypothesis^10,11,12^, forgetting occurs when a memory has a moderate degree of activation (vs. too high or too low) following the instruction to forget. This highlights the contribution of an automatic memory weakening mechanism to deliberate forgetting, and it suggests an alternative strategy for successful forgetting: to weaken an unwanted memory, raise (rather than suppress) its level of activation.

## METHODS

### Subjects

Twenty-four healthy subjects between the ages of 18 and 35 we recruited from the UT Austin student body as well as the surrounding community in accordance with the University of Texas Institutional Review Board. Subjects were compensated at $20 an hour. All subjects were right-handed, had normal or corrected-to-normal vision. Exclusionary criteria included psychiatric disorder, substance abuse and use of psychotropic medication. During the data collection phase, two subjects were excluded for sleeping in the scanner and one additional subject was excluded for claustrophobia. One additional subject was excluded due to data storage malfunction. One final subject was excluded for behavioral performance > 1.5 standard deviations from average performance. A total of 20 subjects (10 female, mean = 23.6 yrs) are included in the reported analyses. An fMRI response box malfunction affected behavioral data recorded for four subjects. As a consequence, in the localizer task, two subjects were not included in the analysis of response latency and accuracy while two others included only accuracy information. For the encoding phase, two subjects were not included in the analysis of encoding task accuracy and response latency, while a third subject contributed only task accuracy information. Critically, these three subjects completed the task and contributed recognition memory task data and were included in the main analyses.

### Stimulus Materials

Experimental materials comprised of colored pictures of scenes, faces and objects. Face stimuli were drawn from a previously published experiment ^19^ and their sources (including www.mac-brain.org/recources.htm). Faces were cropped from the neck down and shown over a white background. A subset of these faces was chosen based on moderate memorability ratings (2.33-4.10, mean: 3.17) from a stimulus evaluation experiment conducted through Amazon.com’s Mechanical Turk^13^. A subset of scenes from the Fine-Grained Image Memorability (FIGRM) Dataset^39^ were used in the present experiment. Scenes were chosen by taking images comprising moderate memorability ratings (2.28–4.38, mean: 3.278, scaled from 1-5) for the task. Objects were drawn from various online sources including Google Images, cropped to exclude any original background, and displayed over a white background. All items were sized to 300 × 300 pixels, and presented using Psychophysics Toolbox Version 3 (PTB-3) in MATLAB 2014a running on an Apple MacBook Air computer running OS X 10.5.

### Procedural Details

Each subject completed three phases for each experimental session (Fig 1). In the localizer phase, subjects performed a perceptual localizer task to train fMRI pattern classifiers on categories of scenes, faces, objects and rest. During the localizer, subjects performed a 1-back task with mini-blocks of items from the three categories of pictures. For the rest category, 18 sec of continuous fixation served as the ‘rest’ mini-block condition. Otherwise, a mini-block consisted of 9 items from the same category shown in succession with 8 sec in between mini-blocks. For each mini-block, 1 or 2 items repeated, thus there was a total of 7-8 unique items per mini-block. Subjects were required to respond ‘not a repeat’ with their right index finger button or ‘repeat’ with their right middle finger button for each item. Within the mini-blocks, each trial began with the presentation of a single item for 1.5 sec, followed by a three horizontally aligned fixation crosses for 50 msec. The localizer phase consisted of 3 localizer runs. Each run included 4 blocks, each block included 1 mini-block of each category type. Across all three runs, the localizer included 90 faces, 90 scenes, 90 objects and 12 mini-blocks of rest. The entire localizer phase lasted approximately 15 min.

The second phase comprised the encoding phase of an item-method directed forgetting task. In this task, subjects were shown either a face or a scene for 3 sec. During the presentation of each item, subjects were instructed to give a subcategory judgment. If a face was presented, subjects were to indicate whether the face was male or female. If a scene was presented, subjects were to indicate whether the scene was of an indoor or outdoor scene. Following the presentation of the item, an instruction cue was given for 6 sec in the form of a yellow fixation cross (‘forget’) or black fixation cross (‘remember’). Subjects were instructed to apply the instruction represented by the cue to the preceding item. Notably, subjects were not encouraged to use any particular strategy, but rather to simply forget or remember the previously presented item. Importantly, critical forget trials were always preceded with a remember trial of the opposing category (e.g., if a face was presented on a forget trial, a scene preceded it in a remember trial) so that our category-specific pattern classification analyses could distinguish trial-specific memory processing (see MVPA section below). In order to discourage anticipation of the forget instruction, we included 60 additional remember trials that were distributed across the experiment to precede other remember trials. Therefore, there were instances in which a remember trial was followed by another remember trial, but a forget trial was never followed by another forget trial. The directed forgetting phase consisted of 6 study runs. Each run included 21 faces and 21 scenes. Across all 6 runs, this phase included 126 faces and 126 scenes, and lasted approximately 38 min.

The third phase of the experiment was a self-paced recognition test conducted outside the scanner. Subjects performed a recognition memory task on a large set of 504 items that included 252 items from the study task (half faces, half scenes) and 252 new items. Subjects were asked to give confidence judgments (‘definitely old, probably old, probably new or definitely new’) to each item presented at test. In order to encourage recognition responses that reflect actual memory of the items, and to discourage responses that reflected the instruction cue given at study (e.g., to discourage a ‘definitely old’ response being given to an item for which the subject remembers being told to forget), confidence responses were assigned points. Subjects were informed of the point system, instructed to maximize their points, and the total point sum was reported to the subject at the end of the test. The point system was as follows: For each old item, a new response (‘probably new’ or ‘definitely new’) was penalized with −1 point while an old response (‘probably old’ or ‘definitely old’) was awarded +1 point. For each new item, an old response (‘probably old’ or ‘definitely old’) was penalized −1 point while a new response (‘probably new’ or ‘definitely new’) was awarded +1 point. Practice items for both localizer and study items were administered prior to the scan session. Test items were not practiced.

### Receiver Operating Characteristic (ROC) curve construction and analysis

Receiver Operating Characteristic (ROC) curve analysis provides a quantitative comparison of memory sensitivity^40^. ROC curve analysis compares memory sensitivity by plotting the proportion of hits (saying an item is old when it was seen before) against the proportion of false alarms (FA, saying an item is old when it was not seen before) over a range of response thresholds. Here we used memory confidence in place of response thresholds^41^. Remember and Forget ROC curves were constructed by plotting hit rate/FA rate pairings from most to least confident, building in a cumulative fashion for each condition. Importantly, as hits rates increase, FA rates also increase – but a more rapid rise in hit rate compared to FA rate in turn, increases the area under the curve produced by the ROC. Greater area below a recognition memory ROC curve describes increased memory sensitivity. We calculated and compared the area under each curve for both Forget and Remember ROC curves.

### fMRI data acquisition

Functional and anatomical MRI data were acquired on a 3T Siemens MRI Scanner (Magnetom Skyra, Siemens AG, Germany) equipped with a 32-channel parallel imaging head coil. Functional scans were acquired with a T2* weighted echo-planar image (EPI) sequence with the following parameters: (TR=1s TE=30 ms, flip angle= 63°, 2.4 mm slices, no gap, 110×110 matrix, FOV=230mm, 56 oblique axial slices, multiband acceleration factor = 4). Slices were acquired in interleaved order. Automatic high order shim was used to orient acquisition parallel to the AC-PC line for full coverage of the brain with limited coverage of the cerebellum. Data was acquired for both localizer and study phases while the test phase was acquired outside the scanner. High-resolution T1-weighted anatomical images were acquired for all subjects using a 3D magnetization-prepared rapid gradient echo (MP-RAGE) pulse sequence (TR = 1.9s, TE = 2.43ms, flip angle = 9°, FOV = 256mm, matrix size 256×256, voxel size 1mm^3^, 192 slices, sagittal acquisition).

### fMRI data analyses

#### Univariate analyses

Functional EPI images were preprocessed and analyzed using SPM12, (http://www.fil.ion.ucl.ac.uk/spm/) implemented under MATLAB R2014a. EPI images were spatially aligned to the mean volume and reoriented parallel to the anterior to posterior commissure plane prior to normalization. All volumes were normalized to the MNI (Montreal Neurological Institute) template EPI* brain and further smoothed 6mm FWHM. We implemented a mass univariate, general linear model analysis primarily to confirm the presence of directed forgetting effects found in previous experiments that used item-method directed forgetting paradigm^18,19.^ We implemented a 2 stage mixed-effects model by first convolving the onset of each Remember and Forget instruction with a canonical hemodynamic response function (HRF) with its temporal and dispersion derivatives. In the first stage, we used the subsequent memory procedure to sort trials from study into items subsequently forgotten or subsequently remembered. Further, we segregated these items into those previously presented with a ‘to-be-forgotten’ instruction or a ‘to-be-remembered’ instruction. In the second stage. we carried these effects of interest forward into a random-effects analysis. We were interested in two primary comparisons: 1) <u>successful forgetting effects</u> – regions demonstrating greater activity for subsequently forgotten, ‘to-be-forgotten’ items than subsequently remembered, ‘to-be-remembered’ items. 2) <u>successful remembering effects</u> - regions demonstrating greater activity for subsequently remembered, ‘to-be-remembered’ items than subsequently forgotten, ‘to-be-forgotten’ items. All effects are reported at an uncorrected threshold of *P* <.001(one sided) with a 20-voxel cluster extent threshold unless otherwise specified.

### Multivariate Pattern Analysis (MVPA)

For MVPA^20^, functional EPI images were preprocessed and analyzed using FSL 5.0 (https://fsl.fmrib.ox.ac.uk/fsl/fslwiki/)^42,43^ subroutines implemented under MATLAB R2014a. Functional images were realigned to the middle volume of the middle (fifth overall) run to correct for motion, and high-pass filtered (128s) to eliminate slow drift. All MVPA analyses were done in native space for each participant (using the Princeton MVPA toolbox (https://github.com/PrincetonUniversity/princeton-mvpa-toolbox) and custom code in MATLAB R2014a).

We used MVPA to quantify the degree of face and scene category-specific neural activity associated with items on forget trials and remember trials. In order to ensure accurate decoding of face and scene categories, we trained a L2–penalized logistic regression classifier (with a penalty (lambda) of 50) on faces, scenes, objects and rest-related activity from the category localizer task. For each mini-block, we trained and tested the classifier on the preprocessed BOLD data from the 18 TRs after the onset of the first item. We shifted regressors by 5 sec to account for hemodynamic delay. Classifier training consisted of using the “leave-one-run-out” method on the 3 localizer runs in which the classifier trains on one run, and tests on the two others, rotating through until all runs are tested. Fig. 1b shows the mean classifier evidence for each category, showing that the classifiers have sufficient sensitivity to discriminate each category of interest.

To decode the directed forgetting task for each participant, we trained classifiers on all localizer data (separately for each participant) and applied them to each TR of the remember trials and forget trials. Here, we produced classification evidence output scores for each 1-sec TR after the trial onset (uncorrected for hemodynamic delay). From the decoded classifier evidence at each time point in each trial, we calculated ‘target’ and ‘nontarget’ evidence by appropriately relabeling the data (e.g., ‘face’ evidence became ‘target’ evidence, and ‘scene’ evidence became ‘nontarget’ evidence on a face trial). Finally, we calculated the differences between target and nontarget evidence to reflect the relative balance of trial-relevant and trial-irrelevant processing at every time point.

Critically, each ‘to-be-forgotten’ item was followed by a ‘to-be-remembered’ item of the opposite category (e.g., a ‘to-be-forgotten’ face followed by a ‘to-be-remembered’ scene). On the other hand, ‘to-be-remembered’ items could be followed by either type of item, including a ‘to-be-remembered’ item from the same category. These trials (38.4% of all remember trials) were not included in any of the analyses to ensure that all items were preceded by an item that was given an opposite instruction (forget or remember) and was drawn from the opposite category (scene or face). While this may seem a considerable proportion of remember trials, it was critical that subjects were unable to anticipate forget trials while during the encoding phase of the trial.

### Regions of interest

All MVPA analyses were conducted within an anatomical ventral temporal mask for each participant. The ventral temporal mask (in MNI space; Montreal Neurological Institute) was defined using boundaries delineated by Grill-Spector and Weiner^42^ and created by merging the temporal fusiform cortex, parahippocampal gyrus, occipital fusiform gyrus and temporal occipital fusiform cortex regions from the Harvard-Oxford atla^43,44,45^ found in FSL 5.0. To create subject-specific masks we co-registered EPI volumes to their own MPRAGE structural volume using FSL FMRIB’s Linear Image Registration Tool (FLIRT)^46,47^. We then used FSL FMRIB’s Non-linear Image Registration Tool (FNIRT) to register structural volumes to MNI space. Individual, native-space ventral temporal masks were created by combining (with the registration parameters for the MPRAGE) and applying a reversed transformation matrix from EPI to MNI stereotaxic space on the ventral temporal mask described above.

### Relating classifier evidence to subsequent memory performance

We utilized the Probabilistic Curve Induction and Testing Toolbox (P-CIT)^11,13^ developed in MATLAB (https://code.google.com/p/p-cit-toolbox) which uses a Bayesian curve-fitting algorithm to estimate the shape of a ‘plasticity curve’ relating neural data (category-specific classifier evidence) and behavioral data (recognition memory confidence scores). In this analysis, we used a ‘super-subject’ procedure in which each participant contributed 96 trials for a total of 1,824 trials for each Forget and Remember instruction condition. We used this fixed-effects analysis because data from each individual subject was insufficient for random effects analysis (see Lewis-Peacock and Norman^13^ for a more detailed explanation) across subjects. Additionally, for this application of P-CIT, we treated pre-instruction (and post-item onset, 1-3s) and post-instruction (4-9s) time intervals as separate events with distinctive processes (perceptual encoding vs. mnemonic processing). This approach to modeling each trial with two neural data points uses the “net effects” procedure^12^ to sum their individual contributions to the single behavioral outcome of remembered or forgotten (see the P-CIT manual for further details). In order to evaluate the reliability of these results, we also implemented a bootstrap resampling procedure^30^ with 1000 iterations.

### Visualization of Results

GLM and GLM-related surface results are visualized the SPM12 canonical render. All subcortical findings are visualized over an MPRAGE volume that is comprised of averaging the MPRAGE volumes specific to this dataset.

### Data and Code Availability

The data and software code used to support the findings of this study are available from the corresponding upon reasonable request.

## Acknowledgements

We thank Morgan Harnois, Katy Hebel and Ellen Crowe for assistance in compiling stimuli and collecting data. We also thank the UT Austin Imaging Research Center, especially Dr. Jeffrey Luci, Chad Cumba and Dr. Donald Nolting and for facility and data collection support. THW was supported by an NIH NRSA post-doctoral training grant (F32NS096962) for this study.

## Contributions

THW, KP, and JALP designed the experiment. THW and KP collected the data. THW and JALP analyzed the data and wrote the paper.

