## Supplementary Materials for "More is less: increased processing of unwanted memories facilitates forgetting"

Supplementary data

Supplementary Table 1. GLM regions identified during item-method intentional forgetting by study memory instruction and subsequent memory outcome. A) Regions demonstrating greater activity for subsequently forgotten items post forget-cue than subsequently remembered items post remember-cue with a threshold of p<.001 and cluster size of 20. B) Regions demonstrating greater activity for subsequently remembered items post remember-cue than subsequently forgotten items post forget-cue with a threshold of p<.005 and a cluster size of 20.

| A. Successful Forgetting | | | | | | | | |
| --- | --- | --- | --- | --- | --- | --- | --- | --- |
| **Coordinates** | | | | | **Cluster size** | **z-value** | | **Approximate Location** |
| 49 | -38 | | | 50 | 661 | 5.26 | | R lateral Parietal |
| 25 | 10 | | | 55 | 1062 | 5.03 | | R dorsolateral prefrontal cortex |
| 6 | -23 | | | 48 | 76 | 4.73 | | posterior cingulate |
| 40 | 13 | | | 2 | 370 | 4.59 | | precuneus |
| 6 | -26 | | | 31 | 172 | 4.14 | |  |
| 8 | 22 | | | 36 | 159 | 4.13 | |  |
| 59 | -28 | | | 10 | 31 | 4.07 | |  |
| -28 | 39 | | | 31 | 71 | 3.99 | |  |
| 20 | 3 | | | 0 | 44 | 3.87 | |  |
| -37 | -52 | | | -48 | 180 | 3.82 | |  |
| 11 | -71 | | | 16 | 93 | 3.62 | |  |
| -11 | 15 | | | 52 | 21 | 3.56 | |  |
| 52 | 8 | | | 16 | 213 | 3.47 | |  |
| B. Successful Remembering | | | | | | | | |
| **Coordinates** | | | | | **Cluster size** | **z-value** | | **location** |
| -30 | | -40 | 7 | | 31 | | 3.48 |  |
| -25 | | -16 | -20 | | 53 | | 3.46 | L hippocampus |
| 16 | | -40 | 19 | | 51 | | 3.44 |  |
| -13 | | 42 | -17 | | 79 | | 3.38 |  |
| -23 | | -38 | 21 | | 24 | | 3.18 |  |

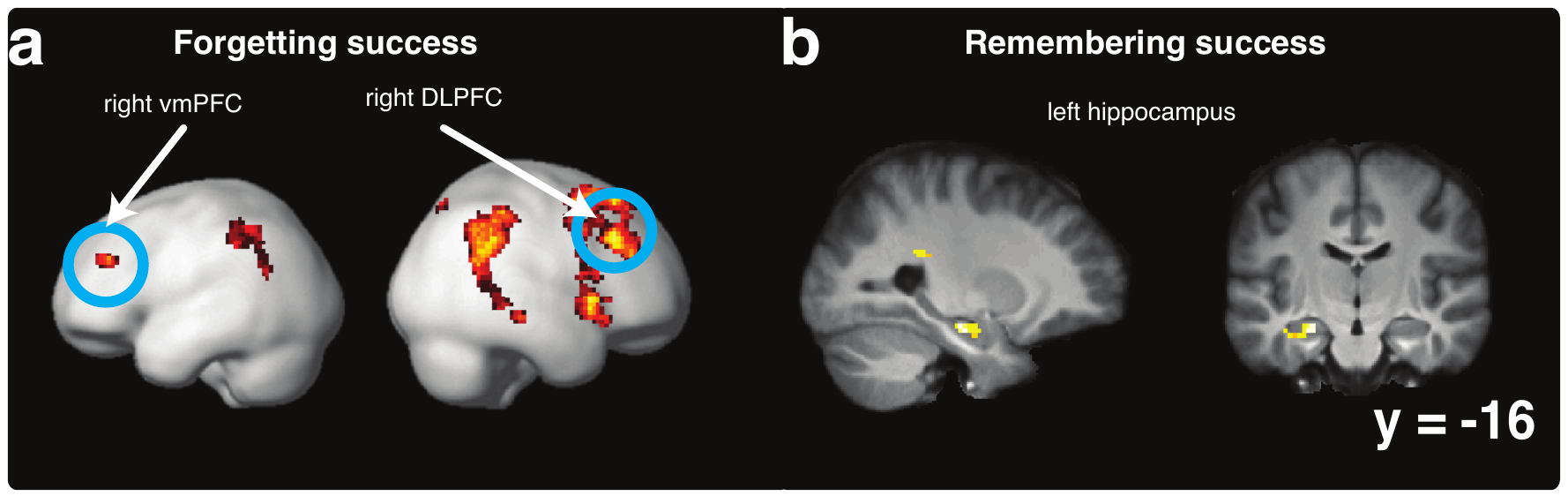
 **Supplementary Figure 1**: Univariate results. a(left): GLM results for forgetting success (greater activity for successful intentional forgetting relative to successful intentional remembering, p<.001, k = 20). Significant clusters include DLPFC (25 10 55; 42 30 36), vmPFC (-28 39 31), posterior cingulate (6 -26 31) and precuneus (-4 -66 50) b: GLM results for remembering success (greater activity for successful intentional remembering relative to successful intentional forgetting, p<.005, k = 20). Significant clusters include left hippocampus (-25 -16 -20).

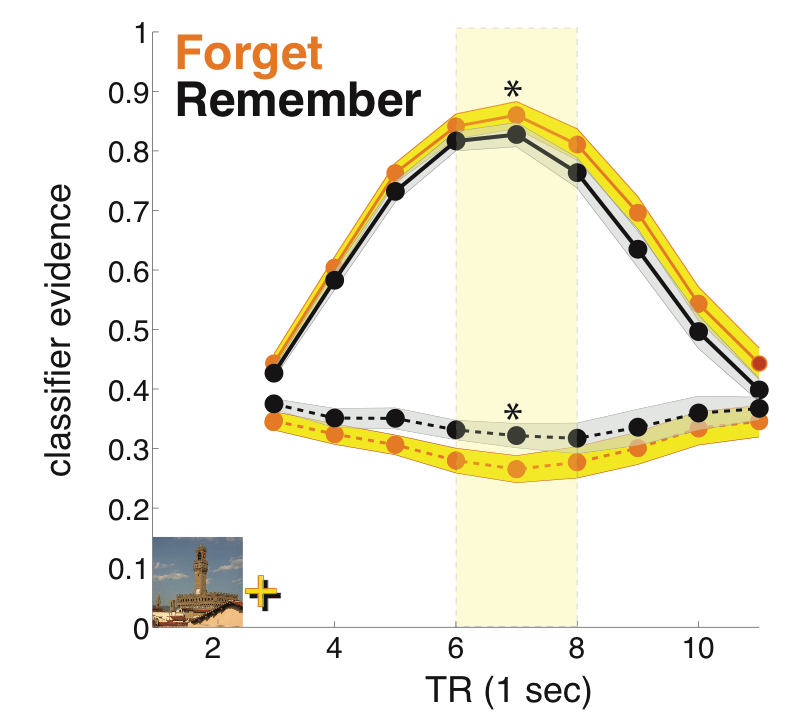

**Supplementary Figure 2.** **Pattern classification of target and non-target processing during the intentional forgetting task**. **(a)** Target and non-target classifier evidence for Forget (yellow) and Remember (black) trials. Classifier evidence scores were not shifted to account for hemodynamic lag. Solid lines represent mean target evidence. Dashed lines represents non-target evidence (Ribbon thickness indicates s.e.m. across participants, n=20). Target evidence 6-8s post item onset for Forget was higher than Remember trials (p<.001) and Non-target evidence 6-8s post item onset for Remember was higher than Forget trials (p<.001).
